# A General Statistic to Test an Optimally Weighted Combination of Common and/or Rare Variants

**DOI:** 10.1101/572115

**Authors:** Jianjun Zhang, Baolin Wu, Qiuying Sha, Shuanglin Zhang, Xuexia Wang

## Abstract

Both genome-wide association study and next generation sequencing data analyses are widely employed in order to identify disease susceptible common and/or rare genetic variants in many large scale genetic studies. Rare variants generally have large effects though they are hard to detect due to their low frequency. Currently, many existing statistical methods for rare variants association studies employ a weighted combination scheme, which usually puts subjective weights or suboptimal weights based on some ad hoc assumptions (e.g. ignoring dependence between rare variants). In this study, we analytically derive optimal weights for both common and rare variants and propose a General and novel approach to Test association between an Optimally Weighted combination of variants (G-TOW) in a gene or pathway for a continuous or dichotomous trait while easily adjusting for covariates. We conduct extensive simulation studies to evaluate the performance of G-TOW. Results of the simulation studies show that G-TOW has properly controlled type I error rates and it is the most powerful test among the methods we compared, when testing effects of either both rare and common variants or rare variants only. We also illustrate the effectiveness of G-TOW using the Genetic Analysis Workshop 17 (GAW17) data. In addition, we applied G-TOW and other competitive methods to test association for schizophrenia. The G-TOW have successfully verified genes FYN and VPS39 which are associated with schizophrenia reported in existing publications. Both of these genes are missed by the weighted sum statistic (WSS) and the sequence kernel association test (SKAT). G-TOW also showed much stronger significance (p-value=0.0037) than our previously developed method named Testing the effect of an Optimally Weighted combination of variants (TOW) (p-value=0.0143) on gene FYN. FYN is a member of the protein-tyrosine kinase oncogene family that phosphorylates glutamate metabotropic receptors and ionotropic N-methyl-d-aspartate (NMDA) receptors. NMDA modulates trafficking, subcellular distribution and function. It is involved in neuronal apoptosis, brain development and synaptic transmission and lower expression, which has been observed in the platelets of schizophrenic patients compared with controls. The application for schizophrenia indicates that G-TOW is a powerful tool in genome-wide association studies.

## Introduction

Genome-wide association studies (GWAS) have identified numerous common variants that are associated with complex human traits (Heid et al., 2010; Allen et al. 2010; Plenge et al., 2007; Saxena et al., 2007; Thomson et al., 2007; Zeggini et al., 2007). However, common variants that have been identified through GWAS can only explain small fractions of estimated trait heritabilities (Bansal et al., 2010; McCarthy et al., 2008; Schork et al., 2009). Rare variants may play an important role in studying the etiology of complex human diseases. Next-generation sequencing technology (Cirulli and Goldstein, 2010) provides potential opportunities to find the missing heritability due to rare genetic variants (Cohen et al., 2006; Ji et al., 2008; Manolio et al., 2009; Marini et al., 2008; Nejentsev et al., 2009; Zhu et al., 2010) since the new sequencing technology makes directly testing for rare variants possible (Andrés et al., 2007).

Due to allelic heterogeneity and the extreme rarity of individual variants, statistical methods used to detect common variants may not be successfully applied to detect rare variants unless sample sizes or effect sizes are very large (Li and Leal, 2008). Recently, several statistical methods for detecting associations of rare variants have been developed. Most of these methods can be divided into two classes: burden tests and quadratic tests. Burden tests collapse rare variants in a genomic region into a single burden variable by using weighted combination and then testing the association between a phenotype and the single burden variable. Choosing appropriate weights is critical to the performance of the burden tests. Different weights reflect different burden test methods, such as the cohort allelic sums test (CAST) (Morgenthaler and Thilly, 2007), the combined multivariate and collapsing (CMC) method (Li and Leal, 2008), the weighted sum statistic (WSS) (Madsen and Browning, 2009), and the variable threshold (VT) method (Price et al., 2010). However, burden tests could suffer loss of statistical power if different directions of effects exist at causal variants. Quadratic tests include tests with statistics of quadratic forms of the score vector such as the sequence kernel association test (SKAT) (Wu et al., 2011), data-adaptive sum (aSUM) (Han and Pan, 2010), adaptive weighting (AW) methods (Sha, Wang, and Zhang, 2013) and weighted sum of squared score (SSU) method (Pan 2009). Even though quadratic tests are robust to the directions of the effects of causal variants, they could be less powerful due to the large number of degrees of freedom of the test statistics when we consider large amounts of rare variants, because the degree of freedom of the quadratic tests depend on the number of rare variants where the quadratic tests usually follows a chi-square distribution with degree freedom related to the number of variants in the considered region.

Sha et al., (2012) propose a method to detect disease associated rare variants by testing an optimally weighted combination of variants (TOW), in which they derived optimal weights by maximizing the score test statistic. From the perspective of the model assumption of the TOW method, which tries to test the effect of a weighted combination of rare variants. The only difference between the traditional burden tests and TOW is that the TOW method chooses optimal weights under a certain criterion, whereas the traditional burden test methods give a subjective weight directly. On the other hand, TOW can also be seen as a quadratic test from the form of the test statistic. Sha et al., (2012) points out that TOW is related to the sequence kernel association test (SKAT) proposed by Wu et al., (2011) and the weighted sum of squared score (SSU) proposed by Pan (2009), which are both quadratic tests. This property means that the TOW method can absorb the advantages of both burden tests and quadratic tests, making it robust to the directions of effects of causal variants and the large number of rare variants.

Due to low minor allele counts of rare causal variants, rare variants are often assumed to be independent either implicitly or explicitly in the development of association test methods. The TOW method is derived under the independent assumption among rare variants. However, simulations based on coalescent models have shown that two variants with low MAFs are likely to be negatively correlated with a negative linkage disequilibrium (LD) value (Hudson, 1985), and ignoring such correlations can reduce the power of aggregation based rare variant association tests (Kinnamon, Hershberger, and Martin, 2012). Talluri and Shete (2013) propose a method for detecting rare variants that utilizes the information of LD between variants (measured by *R*^2^) to select the best subset of rare variants, subsequently improving the power of detecting association. Turkmen and Lin (2017) show that two commonly used LD measures are not capable of detecting LD when rare variants are involved and proposed a new LD measure method.

In this paper, we develop a General and novel approach to Test association between an Optimally Weighted combination of variants (G-TOW) between common and/or rare variants in a gene or pathway and a complex trait of interest. The optimal weights are analytically derived and can be estimated from sampled genotypes and phenotypes under a certain criterion while we consider the correlations among variants in the considered region simultaneously. G-TOW can be applied to both quantitative and qualitative traits, allows covariates, and is robust to directions of effects of causal variants. Extensive simulation studies and applications to the Genetic Analysis Workshop 17 (GAW17) data are used to compare the performance of G-TOW with that of three existing methods (TOW, WSS, and SKAT). Results show that G-TOW is a valid method and consistently more powerful than the other three methods. In addition, we apply all the four methods to a verification study for schizophrenia. G-TOW verifies more important schizophrenia associated genes than other methods.

## Dependence between genetic variants

The correlation between two SNPs are often denoted as LD, which plays a fundamental role in genetic association studies that map genes associated with specific traits of interests. GWAS often rely on high LDs to identify markers that are associated with traits where identified common variants are often in LD with causal variants. However, due to low minor allele frequencies, rare variants are often assumed to be independent. Understanding precisely the LD structure in the human genome including both rare and common variants may lead to better-formed statistical tests that can improve the power of association studies.

Two commonly used statistical measures have been proposed to quantify the degree of LD between a pair of variants: *D*′ and *R*^2^, which have different properties and may be applied for different application purposes (Hill and Robertson, 1968; Lewontin, 1964). Specifically, consider two loci, locus A with minor allele A and locus B with minor allele B. Then *D*′ and *R*^2^ are defined as:

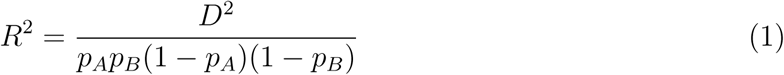

and

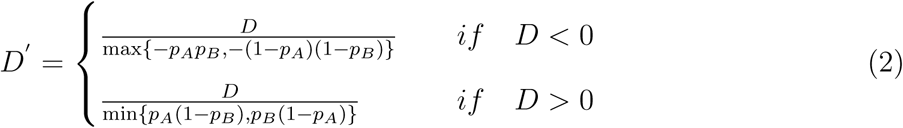

where *D* = *p*_*AB*_ − *p*_*A*_*p*_*B*_ and *p*_*A*_, *p*_*B*_, *p*_*AB*_ denote the frequencies of allele *A*, allele *B*, and haplotype *AB*, respectively. Hudson (1985) had shown that two low frequency variants are likely to be negatively correlated by simulations based on coalescent models, that is *D* < 0 for two rare variants. If locus A and locus B are both rare variants, we can see that the numerator and denominator are approximated by (*p*_*A*_*p*_*B*_)^2^ and *p*_*A*_*p*_*B*_ in equation (1), respectively, and the numerator and denominator are all approximated by −*p*_*A*_*p*_*B*_ in equation (2). This means that regardless of whether these two rare variants are related to each other, the value of *R*^2^ always tends to zero and the value of *D*′ tends to 1. Turkmen and Lin (2017) show that the results from these two measurements are especially sensitive to the selection of tolerance cutoffs for pairs with small MAFs, where the tolerance cutoffs are set in the optimization function to obtain the maximum likelihood estimate of frequencies with the genetic package (Warnes 2013). They also show that *D*\* = sign(*D*)*D*′ will likely attenuate the boundary value to −1, while 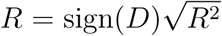 will likely be close to 0 for rare variant pairs even if they come from similar (small) MAFs.

From the aforementioned, we can see that two commonly used LD measures are not capable of quantifying LD when rare variants are involved. The limitations of these two traditional LD measurements lead us to assume that rare variants are independent. Turkmen and Lin (2017) proposed an alternative measure of LD method, the polychoric correlation (PCor), and showed that rare variants in close physical proximity can show strong patterns of local correlation/LD whenever such variants are located on just a few haplotypes. Dickson et al., (2010) showed that some of the signals that have been detected from common variants could come from the effect of rare variants. Correctly handling existing LD among rare variants or between common and rare variants can help researchers to understand the genetic architecture of the human genome and improve power for detecting causal genetic variants. In this study, we propose a novel genetic association test in which we fully consider the correlation situation as either dependent or independent between variants.

## Method

Consider a sample of n individuals and each individual has been genotyped at M variants in a genomic region. Denote *y*_*i*_ as the trait value of the *i*^*th*^ individual for either a quantitative trait or a qualitative trait (1 for cases and 0 for controls for a qualitative trait) and denote *X*_*i*_ = (*x*_*i*1_, *…*, *x*_*iM*_)′ as the genotypic score of the ith individual, where *x*_*im*_ ∈ {0, 1, 2} is the number of minor alleles that the *i*^*th*^ individual has at the *m*^*th*^ variant.

Without loss of generality, we first describe our methods without considering covariates and then extend it to incorporate covariates naturally following Sha et al., (2011). We use the generalized linear model to model the relationship between trait values and genotypes (Nelder and Wedderburn, 1972):

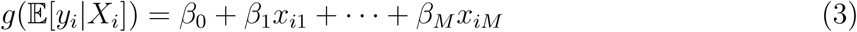

where *g*(·) is a monotone link function and ***β*** = (*β*_0_, *…*, *β*_*M*_) are parameters. Two commonly used models under the generalized linear model framework are the linear model with the identity link for continuous or quantitative traits and the logistic regression model with the Logit link for a binary trait. Under the generalized linear model, the score test statistic to test the null hypothesis *H*_0_ : ***β*** = 0 is given by Sha et al., (2011):

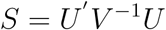

where 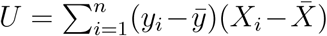 and 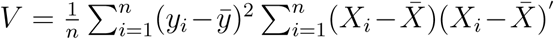. The statistic *S* asymptotically follows a chi-square distribution with *k* = rank(*V*) degrees of freedom (df). However, for rare variants, the score test may suffer power loss due to the sparse data and a large df. To solve this issue, it is general to develop methods for genetic association studies by testing the effect of a weighted combination of variants, 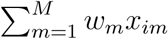, where *w*_*i*_ are the weights and will be given later. That is, the model is written as:

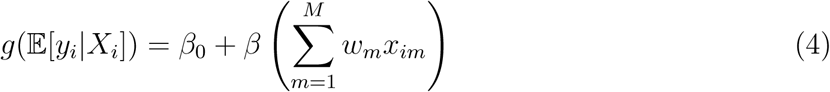

Under this model, it degenerates to test the null hypothesis *H*_0_ : *β* = 0.

Let ***X*** = (*X*_1_, *…*, *X*_*n*_)′ and *Y* = (*y*_1_, *…*, *y*_*n*_)′. To test the effect of the weighted combination of variants 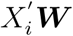 where ***W*** = (*w*_1_, *…*, *w*_*M*_)′, let 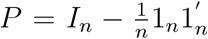 where 1_*n*_ represents a column vector containing all ones, then the score test statistic *S* as a function of *W* can be written as:

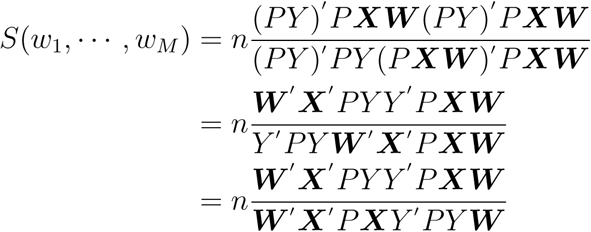

where the second equation holds because we have *P* = *P*′ and *PP*′ = *P* and the third equation holds because *Y*′ *PY* is a constant.

When *D* = ***X***′ *P* ***X*** is positive definite, maximizing *S*(*w*_1_, *…*, *w*_*M*_) is equivalent to maximizing

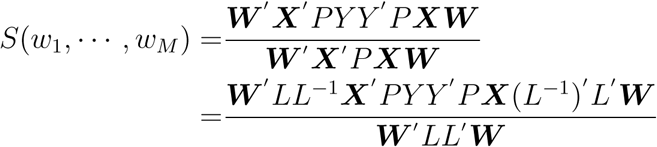

where *L* is the lower triangular matrix obtained from the Cholesky decomposition of *D* = *LL*′. However, when the matrix of *D* is not full rank, it may cause the matrix *D* to be not positive definite (semi-positive definite). If the matrix is not positive definite, we consider two options. First, we can perform correlation pruning so that all pairwise correlation between SNPs is less than 1. Second, we can introduce a ridge parameter *λ*_0_, for which we suggest 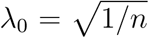 (referred to Baker et al. 2018), where *n* is the number of individuals in the data. This modification, *D* = ***X***′*P* ***X*** + *λ*_0_*I*, allows us to mitigate the flaw of the non-positive matrix *D* in order to avoid the instability. We adopt the first option to in the following simulation study and real data analysis if the matrix is not positive definite.

To maximized *S*(*w*_1_, *…*, *w*_*M*_), let us denote *C* = *L*^−1^***X***′*PY Y*′ *P* ***X***(*L*^−1^)′, and ***c*** be the eigenvector corresponding to the largest eigenvalue of the matrix *C*, then 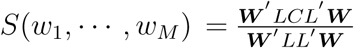 reaches its maximum when *L*′***W*** = ***c***. Therefore, we obtain the optimal weight ***W*** = (*L*′)^−1^***c***. Specially, when we assume that the genetic variants are all independent, the weight of the *m*^*th*^ variant can be expressed as

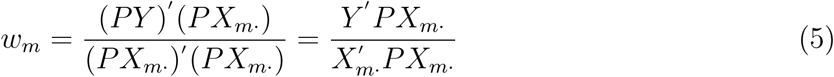

This is the same as the optimal weight in the TOW method (Sha et al., 2012), where *X*_*m·*_ = (*x*_*m*1_, *…*, *x*_*mn*_). The equation (5) is equivalent to 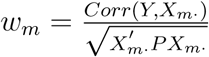 where the numerator is the correlation coefficient between the phenotype *Y* and the *m*^*th*^ variant genotypic score *X*_*m·*_ and the denominator can be viewed as the variance of the *X*_*m*_. This means that *w*_*m*_ has the same direction as the correlation between the phenotype *Y* and the *m*^*th*^ variant *X*_*m*_, and puts big weight to the variant that has strong association with the trait of interest and to the variant with low variance which usually is a low frequency variant.

We define the General statistic to Test the effect of the Optimally Weighted combination (G-TOW) of variants, ***XW***, as:

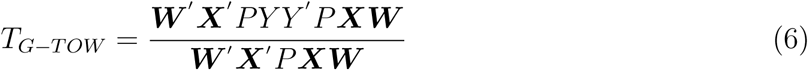

We use permutation methods to evaluate the P-value of *T*_*G-TOW*_. Tow derive the weights to maximize *S*(*w*_1_, *…*, *w*_*M*_) based on the assumption that rare variants are independent. As we mentioned above, correlations between variants play an important role for association analysis and ignoring such correlations can reduce the power of association tests, especially those low frequent variants based association tests.

The G-TOW method can be extended to incorporate covariates. Suppose that there are *p* covariates. Let *z*_*il*_ denote the *l*^*th*^ covariate of the *i*^*th*^ individual. We adjust both trait value *y*_*i*_ and genotypic score *x*_*im*_ for the covariates by using linear regression models. That is,

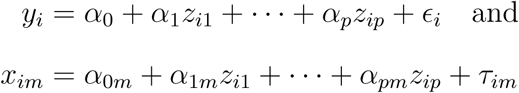

Let 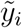 and 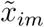 denote the residuals of *y*_*i*_ and *x*_*im*_, respectively. We incorporate the covariate effects in G-TOW by replacing *y*_*i*_ and *x*_*im*_ in equation (6) by 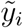 and 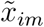. With covariates, the statistic of G-TOW is defined as:

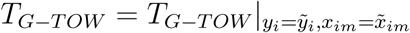

## Comparison of tests

We compared our method, (G-TOW) with three other approaches: 1) the weighted statistic method, (WSS), (Madsen and Browning 2009); 2) Testing the effect of the Optimally Weighted combination of variants, (TOW), (Sha et al., 2012); 3)Sequence Kernel Association Test, (SKAT), (Wu et al., 2011).

## Simulation

Gene TP53 is crucial in multicellular organisms, where it prevents cancer formation, thus, functions as a tumor suppressor. In simulation studies, we generate genotype data using the haplotypes of gene TP53 obtained from the 1000 Genome Project http://www.sph.umich.edu/csg/abecasis/MACH/download/1000G-2010-08.html. This dataset contains 566 hyplotypes for 74 variants (27 common variants and 47 rare variants). Here, we divide variants into rare (MAF<the rare variant threshold (RVT)) and common (MAF*>*RVT), and consider RVT as 0.05. In our simulation studies, we generate genotypes for individuals based on this haplotype pool. To generate the genotype of an individual, we select two haplotypes according to the haplotype frequencies.

### Correlation analysis of the gene TP53

To be able to empirically illustrate the effects of varying values of MAF on correlation measures, we classified SNPs into five index groups using the following cutoffs: Index 1 (MAF*≥*0.05), Index 2 (0.03≤MAF<0.05), Index 3 (0.02≤ MAF<0.03), Index 4 (0.01≤MAF<0.02) and Index 5 (MAF<0.01). Here, Index 1 represents common variants, whereas the remaining indices represent rare variants. Index 5 depicts the most extreme rare scenario. Detailed information for this classification and the number of SNPs in each index set for the dataset is summarized in Table 1.

**Table 1:** Index Definitions for TP53 Data based on MAF Cutoffs

| Group | MAF Range | #SNPs |
| --- | --- | --- |
| Index 1 | $\text{MAF} \geq 0.05$ | 27 |
| Index 2 | $0.03 \leq \text{MAF} < 0.05$ | 8 |
| Index 3 | $0.02 \leq \text{MAF} < 0.03$ | 4 |
| Index 4 | $0.01 \leq \text{MAF} < 0.02$ | 6 |
| Index 5 | $\text{MAF} < 0.01$ | 29 |

For illustration purpose, we have chosen several SNP pairs from the TP53 Data to demonstrate the specific effects of varying values of MAF on *R*^2^, *D*′, *R, D*\* and PCor (Turkmen and Lin 2017). Table 2 summarizes the results reported with different LD measures and it indicates the limitations of existing LD measures for rare variants. As shown in Table 2, if a SNP is rare, it is very likely that the frequency of the haplotype carrying the minor allele can not be estimated accurately, whereas the two known popular LD measures *R*^2^ and *D*′ are calculated under the assumption that the population distribution of haplotypes is known. This will lead to *D*′ attaining 1 or *R*^2^ tending to 0 regardless of the level of real LD. From the values of PCor, we can see that these correlations/LD exist between rare variants or even rare and common variants in the gene TP53 data.

**Table 2:** *R*^2^, *D*′, *R, D*\* and PCor values for the Selected SNP-Pairs on TP53 Data

| | Index | MAF1 | MAF2 | $R^2$ | $D'$ | R | $D^*$ | PCor |
| --- | --- | --- | --- | --- | --- | --- | --- | --- |
| 1 | 1 | 0.297 | 0.290 | 0.844 | 0.936 | 0.919 | 0.936 | 0.993 |
| 1 | 2 | 0.297 | 0.040 | 0.099 | 1.000 | 0.313 | 1.000 | 0.989 |
| 1 | 3 | 0.297 | 0.025 | 0.060 | 1.000 | 0.246 | 1.000 | 0.983 |
| 1 | 4 | 0.297 | 0.019 | 0.008 | 1.000 | -0.089 | -1.000 | -0.951 |
| 1 | 5 | 0.297 | 0.003 | 0.001 | 1.000 | -0.034 | -1.000 | -0.942 |
| 2 | 2 | 0.040 | 0.042 | 0.950 | 1.000 | 0.974 | 1.000 | 0.999 |
| 2 | 3 | 0.040 | 0.025 | 0.001 | 1.000 | -0.032 | -1.000 | -0.990 |
| 2 | 4 | 0.040 | 0.019 | 0.001 | 1.000 | -0.028 | -1.000 | -0.898 |
| 2 | 5 | 0.040 | 0.003 | 0.000 | 1.000 | -0.010 | -1.000 | -0.894 |
| 3 | 3 | 0.025 | 0.028 | 0.907 | 1.000 | 0.952 | 1.000 | 0.999 |
| 3 | 4 | 0.025 | 0.019 | 0.001 | 1.000 | -0.021 | -1.000 | -0.991 |
| 3 | 5 | 0.025 | 0.003 | 0.000 | 1.000 | -0.008 | -1.000 | -0.890 |
| 4 | 4 | 0.019 | 0.015 | 0.000 | 1.000 | -0.017 | -1.000 | -0.899 |
| 4 | 5 | 0.019 | 0.003 | 0.000 | 1.000 | -0.007 | -1.000 | -0.821 |
| 5 | 5 | 0.003 | 0.001 | 0.000 | 1.000 | -0.001 | -1.000 | -0.820 |

### Simulation setting

To assess the performance of the proposed method G-TOW, we conduct extensive simulation studies based on the gene TP53 data. We compare the performance of G-TOW with these existing methods TOW, WSS, and SKAT.

To evaluate type I error rates of the four methods, we generate trait values which are independent of genotypes by using the following model (7) :

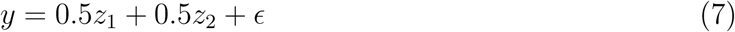

where *z*_1_ is a continuous covariate generated from a standard normal distribution, *z*_2_ is a binary covariate taking values 0 and 1 with a probability of 0.5, and *E* is the error term following a standard normal distribution.

To evaluate power, we randomly choose one common variant and 20% or 40% rare variants (*n*_*c*_) as causal variants where the common variant and half of rare variants (*n*_*r*_ = *n*_*c*_*/*2) are risk variants and the other half of the rare variants (*n*_*p*_ = *n*_*c*_*/*2) are protective variants. Denote 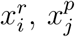 and *x*^*c*^ as genotypes of the *i*^*th*^ risk rare variant, the *j*^*th*^ protective rare variant, and the common variant, respectively. Then we generate a quantitative trait using the following model (8):

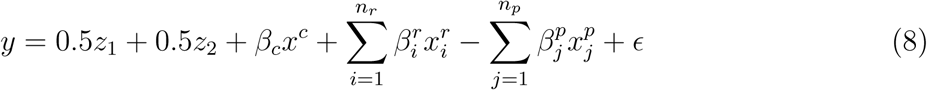

where *z*_1_, *z*_2_ and *ϵ* are the same as those in equation (7). In model (8), 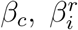 and 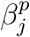 are coefficients. The values of these three coefficients depend on the total heritability *h*_*total*_ and the ratio (*R*^0^) of heritability of the rare causal variant to heritability of the common causal variant. For given *h*_*total*_ and *R*^0^, we can calculate the heritability of the rare casual variants and the common causal variants, respectively. We assume that all the rare causal variants have the same heritability such that rarer variants have larger effects. When we consider common variant and rare variants simultaneously, the values of 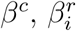 and 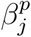 can be determined by:

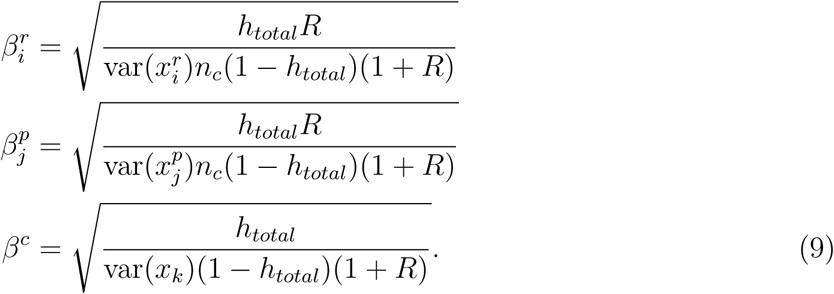

. When we only consider rare variants, the values of 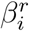 and 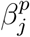 are given as:

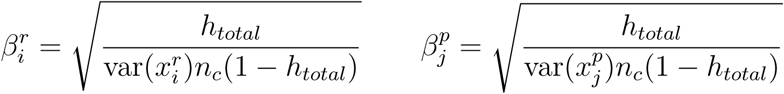

To generate a qualitative trait, we use a liability threshold model based on the quantitative trait (continuous phenotype) described above. An individual is defined to be affected (case) if the individual’s phenotype is at least one standard deviation larger than the phenotypic mean which yields a approximate prevalence of 16% for the simulated disease in a general population.

### Simulation results

For type I error evaluation, we consider different disease models (with or without covariates) for different types of traits (qualitative trait or quantitative trait). The type I error rates are evaluated using 1000 replicated samples and the P-values are estimated using 10,000 permutations. For the 1000 replicated samples, the 95% confidence intervals (CIs) for the estimated type I error rates at nominal levels 0.05, 0.01, and 0.001 are (0.036, 0.064), (0.004, 0.016), and (0.00095, 0.00295), respectively. The estimated type I error rates of these four tests (G-TOW, TOW, WSS and SKAT) are summarized in Table 3. From this table, we can see that all of the estimated type I error rates are either within 95% CIs or close to the bound of the corresponding 95% CIs, which indicate that the existing and proposed methods are valid.

**Table 3:** The estimated type I error rates of the four test methods

| Quantitative trait |  |  |  |  |  |
| --- | --- | --- | --- | --- | --- |
| | α-level | G-TOW | TOW | WSS | SKAT |
| $n = 1000$ | 0.05 | 0.047 | 0.052 | 0.056 | 0.055 |
|  | 0.01 | 0.012 | 0.011 | 0.010 | 0.009 |
|  | 0.001 | 0.000 | 0.001 | 0.001 | 0.000 |
| Qualitative trait |  |  |  |  |  |
| $n = 1000$ | 0.05 | 0.048 | 0.053 | 0.056 | 0.046 |
|  | 0.01 | 0.011 | 0.007 | 0.008 | 0.008 |
|  | 0.001 | 0.001 | 0.001 | 0.000 | 0.001 |

In power comparisons, the P-values of G-TOW, TOW and SKAT are estimated using 1,000 permutations, while the P-values of WSS are estimated by asymptotic distributions. The power of each of the four tests is evaluated using 1,000 replicated samples at the significance level of 0.05 (Figure 1-4). For power comparisons, we consider two different cases: (1) rare causal variants in which all causal variants are rare (MAF<RVT) and (2) both rare and common causal variants in which there is one common variant and the heritability of the common variant is twice as big as the heritability of all the rare causal variants.

**Figure 1:**
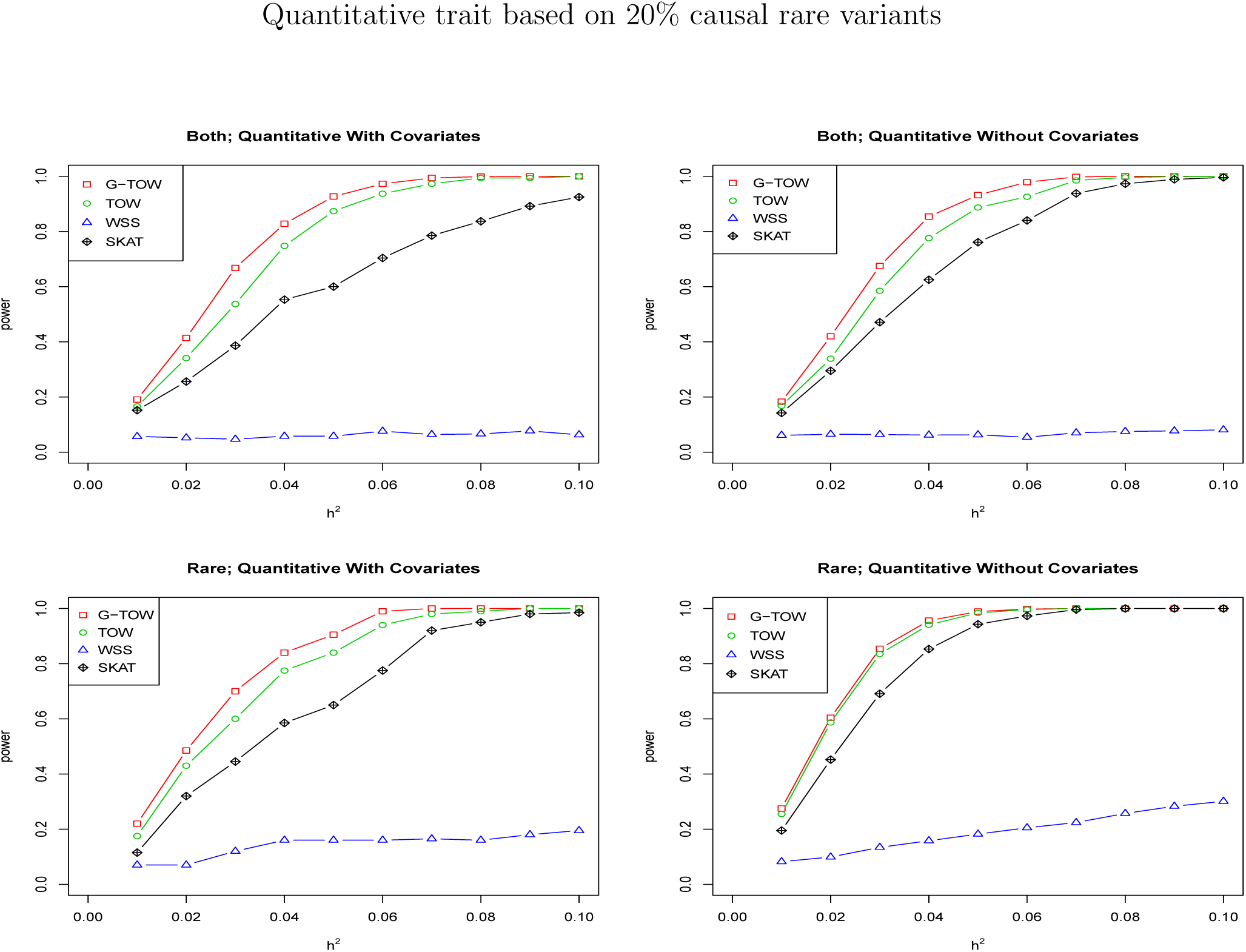
Power comparisons of the four tests (G-TOW, TOW, WSS and SKAT) for the power as a function of heritability for a quantitative trait. Rare means that all causal variants are rare. Both means that causal variants contain both rare and common (one common variant) and the heritability of the common variant is as twice as the heritability of all the rare causal variants. The x-axis represents the total heritability of all causal variants. Sample size is 1,000 and 20% of rare variants are causal, half of rare causal variants are risk and the other half of causal variants are positive. The powers (y-axis) are evaluated at a significance level of 0.05.

Power comparisons of the four tests (G-TOW, TOW, WSS and SKAT) as a function of heritability for a quantitative trait are given in Figure 1 (20% causal rare variants) and Figure 3 (40% causal rare variants). As shown in Figure 1 or Figure 3, G-TOW and TOW are consistently more powerful than WSS and SKAT. We can also see that G-TOW is the most powerful test when the causal variants contain both common and rare variants. G-TOW is the most powerful test when the heritability is less than 0.05 and the casual variants are all rare variants. The power of both G-TOW and TOW is close to 1 when the heritability is greater than 0.05. The method SKAT performs worse than G-TOW and TOW under all of cases in Figure 1 and Figure 3, because the MAFs of a large portion of rare causal variants are under 0.01 and SKAT will lose power when the MAFs of causal variants are not in the range (0.01,0.035) (Yang et al. 2017). G-TOW is more powerful than TOW, especially when the causal variants contain both common and rare variants because TOW doesn’t consider the correlation between variants. The method WSS is the least powerful test, because this method will suffer substantial loss of power when both risk and protective variants are present.

**Figure 2:**
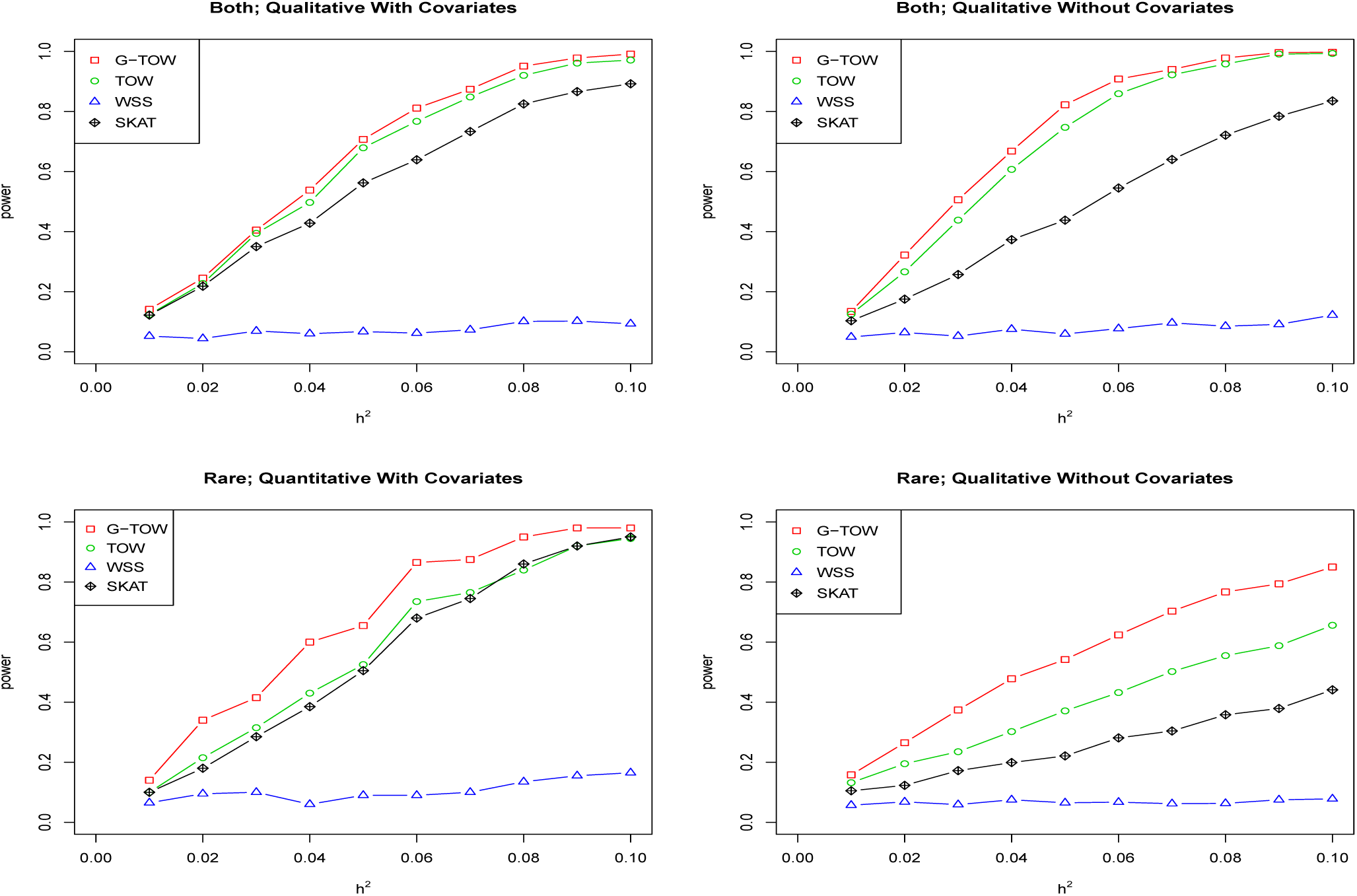
Power comparisons of the four tests (G-TOW, TOW, WSS and SKAT) for the power as a function of heritability for a qualitative trait. Rare means that all causal variants are rare. Both means that causal variants contain both rare and common (one common variant) and the heritability of the common variant is as twice as the heritability of all the rare causal variants. The x-axis represents the total heritability of all causal variants. Sample size is 1,000 and 20% of rare variants are causal, half of rare causal variants are risk and the other half of causal variants are positive. The powers (y-axis) are evaluated at a significance level of 0.05.

**Figure 3:**
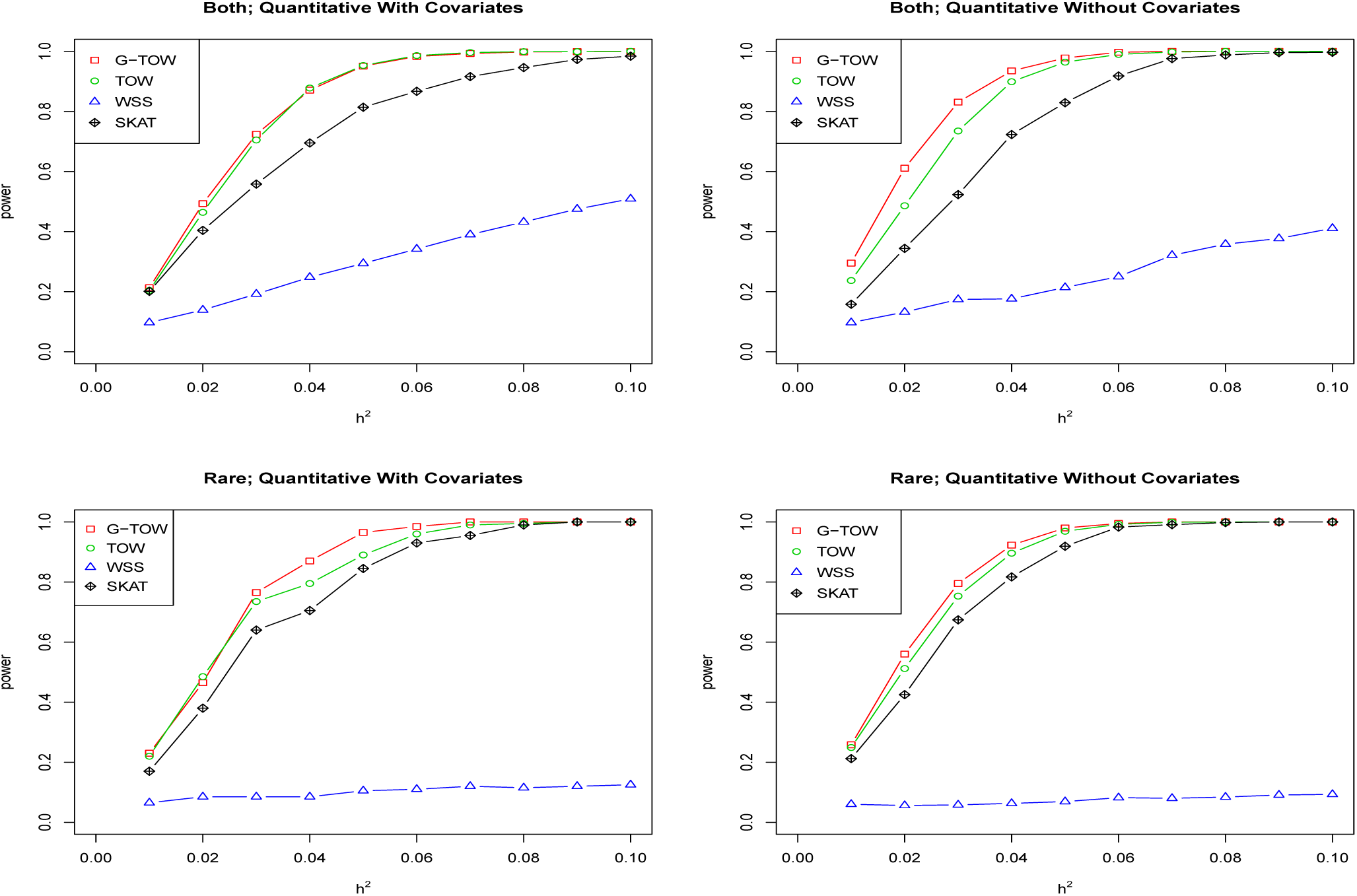
Power comparisons of the four tests (G-TOW, TOW, WSS and SKAT) for the power as a function of heritability for a quantitative trait. Rare means that all causal variants are rare. Both means that causal variants contain both rare and common (one common variant) and the heritability of the common variant is as twice as the heritability of all the rare causal variants. The x-axis represents the total heritability of all causal variants. Sample size is 1,000 and 40% of rare variants are causal, half of rare causal variants are risk and the other half of causal variants are positive. The powers (y-axis) are evaluated at a significance level of 0.05.

Power comparisons of the four tests for different values of heritability based on a qualitative trait are given in Figure 2 (20% causal rare variants) and Figure 4 (40% causal rare variants). Comparing Figure 1 with Figure 2 (or Figure 3 with Figure 4), we can see that the patterns of power comparisons based on a qualitative trait are very similar to that based on a quantitative trait. However, it seems that the power of all four tests in most cases is smaller based on a qualitative trait than based on a quantitative trait. This may because that converting to qualitative variables from quantitative variables results in some information loss. Comparing the case of 20% causal rare variants in Figure 1-2 with that of 40% causal rare variants in Figure 3-4, we can see that patterns of power comparisons based on 20% causal rare variants are very similar to those based on 40% causal rare variants. The power of G-TOW, TOW and SKAT is relatively robust to the increase of neutral variants while the power of WSS still suffers a substantial loss of power in the presence of both risk and protective variants.

**Figure 4:**
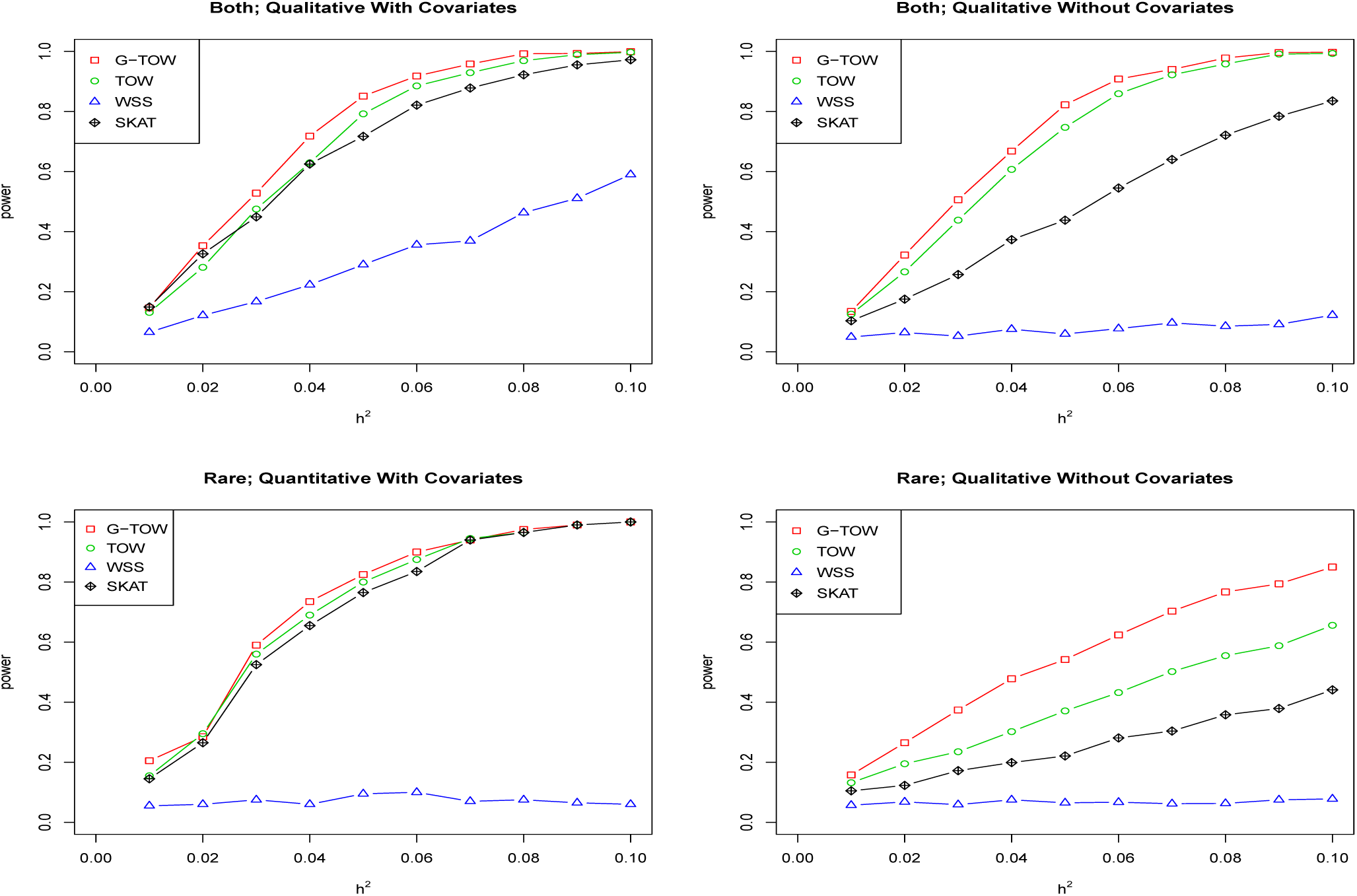
Power comparisons of the four tests (G-TOW, TOW, WSS and SKAT) for the power as a function of heritability for a qualitative trait. Rare means that all causal variants are rare. Both means that causal variants contain both rare and common (one common variant) variants. The heritability of the common variant is as twice as the heritability of all the rare causal variants. The x-axis represents the total heritability of all causal variants. Sample size is 1,000 and 40% of rare variants are causal, half of rare causal variants are risk and the other half of causal variants are positive. The powers (y-axis) are evaluated at a significance level of 0.05.

Because the method TOW can be viewed as a special case of G-TOW, G-TOW is consistently more powerful than TOW when correlation between variants exist and has similar power when the variants are all independent. In summary, G-TOW is consistently more powerful or has similar power to the method TOW, and it is more powerful than WSS and SKAT. The power of G-TOW is also relatively robust to an increase in the percentage of rare variants.

## Analysis of the GAW17 dataset

The GAW17 dataset consists of real genotypes and 200 replicates of the simulated phenotypes for a a collection of 697 unrelated individuals, which is provided by the 1000 Genomes Project for their pilot3 study. A total of three quantitative traits (Q1, Q2, and Q4), are simulated and covariates include age, sex, and smoking status. Because quantitative trait Q4 is not influenced by any of the genotyped exonic SNPs, we do not consider Q4 for the purpose of power comparisons. The P-values of G-TOW, TOW, and SKAT are evaluated by 10,000 permutations and the P-values of WSS are evaluated by asymptotic distributions. For a better comparison of these four test methods, we merge two replicates into one replicate to increase the sample size and all powers are estimated at a significance level of 0.001. In all cases, all causal variants are risk variants. There are nine causal genes for Q1 and thirteen causal genes for Q2. We omit causal genes that have one variant, causal genes in which there is only one copy of haplotype among the individuals for each causal variant, causal genes in which all of the four tests have 100% power, and causal genes in which all of the four tests have a power less than 10%. There are four genes associated with Q1 and seven causal genes with Q2. The powers of WSS, TOW and SKAT are not consistent with those in Table 2 of Sha et al., (2012), where they did not adjust trait values and genotypes for covariates.

The powers of the four tests to detect association between each of the 11 causal genes and Q1 or Q2 are given in Table 4. As shown in Table 2, G-TOW is the most powerful test in four out of 11 genes, WSS is the most powerful test in three out of 11 genes, TOW is the most powerful test in one out of 11 genes and SKAT is the most powerful test in three out of 11 genes. Comparing G-TOW with TOW, G-TOW is either most powerful test or has similar power to the TOW method in ten of 11 casual genes.

**Table 4:** Power of the four methods to test the association between each of the four causal genes and quantitative trait Q1 and between each of the seven causal genes and quantitative trait Q2

| Traits | Gene name | No. of variants, no. | Min, max, | G-TOW | TOW | WSS | SKAT |
| --- | --- | --- | --- | --- | --- | --- | --- |
|  |  | of causal variants | mean MAF |  |  |  |  |
| Q1 | ARNT | 18,5 | 0.07,1.15,0.33 | 0.58 | 0.59 | 0.08 | **0.94** |
|  | FLT4 | 10,2 | 0.07,0.14,0.11 | 0.66 | 0.69 | **0.85** | 0.66 |
|  | HIF1A | 8,4 | 0.07,1.22,0.39 | 0.57 | 0.59 | 0.62 | **0.90** |
|  | VEGFA | 6,1 | 0.22,0.22,0.22 | 0.17 | 0.16 | **0.35** | 0.11 |
| Q2 | BCHE | 29,13 | 0.07,0.29,0.10 | **0.38** | 0.37 | 0.17 | 0.13 |
|  | LPL | 20,3 | 0.07,1.58,0.60 | 0.25 | 0.23 | 0.00 | **0.36** |
|  | PDGFD | 11,4 | 0.07,0.86,0.29 | **0.18** | 0.15 | 0.01 | 0.14 |
|  | SIRT1 | 23,9 | 0.07,0.22,0.12 | 0.31 | **0.61** | 0.34 | 0.56 |
|  | SREBF1 | 24,10 | 0.07,0.43,0.22 | 0.22 | 0.17 | **0.24** | 0.04 |
|  | VNN1 | 7,2 | 0.57,17.1,8.82 | **0.83** | 0.71 | 0.08 | 0.03 |
|  | VNN3 | 15,7 | 0.07,9.83,2.06 | **0.72** | 0.53 | 0.31 | 0.37 |

Note: Min, max, mean MAF: the minimum, maximum, and mean MAF (in percentage) at causal variants. In each row, the boldfaced number represents the highest power in the row.

## Analysis of the Schizophrenia dataset

Schizophrenia is a severe and disabling mental illness with onset typically in early adult life. It is associated with low fecundity but nevertheless remains fairly common with a lifetime prevalence of around 1% (Power et al., 2013). Many genes including common and rare variants have been reported association with schizophrenia (Curtis et al. 2017, Genovese et al. 2016, Purcell et al. 2014, Fromer et al. 2014, Takata et al. 2014, Rujescu et al. 2009, Xu et al. 2012, Millar et al. 2000). In this study, we seek to verify the 14 associated genes (see Table 5) for schizophrenia by analyzing exome-sequencing of 2,536 schizophrenia cases and 2,543 controls (total 5079 individuals) of a Swedish sample. SNPs on the 14 genes with MAF that equals to zero and missing rate greater than 20% are excluded in the analysis.

**Table 5:** P-values of these four methods to test association between genes and case-control trait

| Gene name | Min, max, | G-TOW | TOW | WSS | SKAT |
| --- | --- | --- | --- | --- | --- |
|  | mean MAF |  |  |  |  |
| ARC | 0.0001, 0.4618, 0.0339 | 0.2583 | 0.2817 | 0.4645 | 0.7751 |
| DISC1 | 0.0001, 0.3111, 0.0135 | 0.7645 | 0.6939 | 0.7736 | 0.4124 |
| DISC2 | 0.0001, 0.1579, 0.0309 | 0.5306 | 0.5056 | 0.6880 | 0.5407 |
| DPYD | 0.0001, 0.2481, 0.0102 | 0.1475 | 0.1959 | 0.4782 | 0.1119 |
| FMR1 | 0.0001, 0.0847, 0.0078 | **0.0339** | 0.0598 | **0.0313** | **0.0252** |
| FYN | 0.0001, 0.1377, 0.0064 | ***0.0037*** | **0.0143** | 0.9254 | 0.1628 |
| KL | 0.0001, 0.4038, 0.0213 | 0.5505 | 0.5829 | 0.6533 | 0.8737 |
| LAMA2 | 0.0001, 0.4669, 0.0183 | 0.1154 | **0.0138** | 0.5967 | 0.1166 |
| NMDAR | 0.0001, 0.4118, 0.0192 | 0.3431 | 0.3272 | 0.3804 | 0.4312 |
| NRXN1 | 0.0001, 0.0090, 0.0009 | 0.5764 | 0.4227 | **0.0303** | 0.4997 |
| SETD1A | 0.0001, 0.3788, 0.0037 | 0.6878 | 0.5235 | 0.6386 | 0.7821 |
| TAF13 | 0.0001, 0.1276, 0.0109 | 0.7455 | 0.7414 | 0.3661 | 0.7041 |
| TRRAP | 0.0001, 0.2383, 0.0034 | 0.1046 | 0.4087 | 0.1040 | 0.6638 |
| VPS39 | 0.001, 0.1830, 0.0115 | **0.0442** | **0.0245** | 0.4767 | 0.0752 |

Note: Min, max, mean MAF: the minimum, maximum, and mean MAF (in percentage) at causal variants. In each row, the boldfaced number represents the significant P-value in the row (significant level = 0.05); the italic number in each row represents the nearly significant P-value after correcting for multiple testing issue (significant level = 0.0036).

The P-values of these four methods to test the association between each of the 14 genes and the case-control trait of schizophrenia are given in Table 5. As shown in Table 5, the number of genes verified by these four methods G-TOW, TOW, WSS and SKAT are 3,3,2,1, respectively, when we use the significant level as 0.05. When we consider multiple testing issue by using a significant level as 0.0036, among these four methods, only gene FYN identified by the G-TOW (P-value = 0.0037). At this gene, the P-value of G-TOW is an order of magnitude smaller than the P-value of TOW method. Only TOW method can’t verify FMR1 gene. Compared to the other three methods, we can see that our method G-TOW performs best overall from Table 5. For the three verified genes by G-TOW (using significant level as 0.05), gene FYN, VPS39 and FMR1 have been reported significant association with schizophrenia by Curtis et al. (2017), Xu et al. (2012) and Purcell et al. (2014), respectively. FYN was identified as a “prioritized candidate gene” and an intronic marker, rs7757969, on this gene was significant at *p* = 4.8*10^−8^ (Li et al., 2017) and the gene FYN is involved in neuronal apoptosis, brain development and synaptic transmission and lower expression has been observed in the platelets of schizophrenic patients compared with controls. Xu et al. (2012) had showed that gene VPS39 was affected by recurrent novo events in a sequenced 795 exomes from 231 parent-proband trios enriched for sporadic schizophrenia cases, as well as 34 unaffected trios. Purcell et al. (2014) proposed that targets of the fragile *times* mental retardation protein (FMRP, product of FMR1) were enriched for in cases with schizophrenia.

## Discussion

Most of the recently developed methods for rare variants association studies essentially assume that rare variants are independent. Using the two popular LD measures *D*′ and *R*^2^ is not capable of detecting the correlation involving rare variants. Ignoring such correlation between variants will reduce the power of rare variants association tests. Motivated by TOW method, we analytically derived the optimal weights where we fully considered the correlations between variants no matter common or rare. Based on the optimal weights, we proposed G-TOW to test the effect of the optimally weighted combination of variants. We used extensive simulation studies and application to the GAW17 dataset and the real schizophrenia data to compare the performance of our G-TOW method with existing methods (TOW, WSS and SKAT). The results show that our method is consistently more powerful than the other methods we compared with.

When we use burden tests to test rare variants, most of them put subjective weights on variants (usually put large weights on rare variants and small weights on common variants). By putting wrong weights on variants, these methods lose power when testing the effects of both rare and common variants. To test the effects of both rare and common variants, WSS puts weights as the inverse square root of the expected variance based on allele frequencies on rare and common variants. By putting positive weights on variants, WSS loses power when causal variants contain both risk and protective variants. The SKAT method puts the beta distribution density function with pre-specified parameters determined by MAFs of variants in the data. However, SKAT has better performance only for those variants with MAF in a certain range (Sha et al., 2012). Our proposed G-TOW, choosing weights adaptively by combining correlation between variants under a certain criterion, has relatively better performance than TOW in testing the effects of both rare and common variants or rare variants only, since TOW ignores the correlation between variants.

## Acknowledgments

Q Sha and S Zhang are funded by the National Human Genome Research Institute of the National Institutes of Health under Award Number R15HG008209. The content is solely the responsibility of the authors and does not necessarily represent the official views of the National Institutes of Health. The Genetic Analysis workshops are supported by NIH grant R01 GM031575 from the National Institute of General Medical Sciences. Preparation of the Genetic Analysis Workshop 17 Simulated Exome Dataset was supported in part by NIH R01 MH059490 and used sequencing data from the 1,000 Genomes Project (http://www.1000genomes.org). The Schizophrenia data/analyses presented in the current publication are based on the use of study data downloaded from the dbGaP web site, under dpGaP accession: phs000473.v2.p2. The authors have no conflict of interest to declare.

